# Sequencing of 15,622 gene-bearing BACs reveals new features of the barley genome

**DOI:** 10.1101/018978

**Authors:** María Muñoz-Amatriaín, S Lonardi, MC Luo, K Madishetty, JT Svensson, MJ Moscou, S Wanamaker, T Jiang, A Kleinhofs, GJ Muehlbauer, RP Wise, N Stein, Y Ma, E Rodriguez, D Kudrna, PR Bhat, S Chao, P Condamine, S Heinen, J Resnik, R Wing, HN Witt, M Alpert, M Beccuti, S Bozdag, F Cordero, H Mirebrahim, R Ounit, Y Wu, F You, J Zheng, H Šimková, J Doležel, J Grimwood, J Schmutz, D Duma, L Altschmied, T Blake, P Bregitzer, L Cooper, M Dilbirligi, A Falk, L Feiz, A Graner, P Gustafson, PM Hayes, P Lemaux, J Mammadov, TJ Close

## Abstract

Barley (*Hordeum vulgare* L.) possesses a large and highly repetitive genome of 5.1 Gb that has hindered the development of a complete sequence. In 2012, the International Barley Sequencing Consortium released a resource integrating whole-genome shotgun sequences with a physical and genetic framework. However, since only 6,278 BACs in the physical map were sequenced, detailed fine structure was limited. To gain access to the gene-containing portion of the barley genome at high resolution, we identified and sequenced 15,622 BACs representing the minimal tiling path of 72,052 physical mapped gene-bearing BACs. This generated about 1.7 Gb of genomic sequence containing 17,386 annotated barley genes. Exploration of the sequenced BACs revealed that although distal ends of chromosomes contain most of the gene-enriched BACs and are characterized by high rates of recombination, there are also gene-dense regions with suppressed recombination. Knowledge of these deviant regions is relevant to trait introgression, genome-wide association studies, genomic selection model development and map-based cloning strategies. Sequences and their gene and SNP annotations can be accessed and exported via http://harvest-web.org/hweb/utilmenu.wc or through the software HarvEST:Barley (download from harvest.ucr.edu). In the latter, we have implemented a synteny viewer between barley and *Aegilops tauschii* to aid in comparative genome analysis.

## INTRODUCTION

Since Neolithic times, barley has played a major role as a source of food, feed, and beer (1). The ability of barley to adapt to marginal environments together with the distinctive grain characteristics make barley a versatile crop that is grown worldwide (2). However, genes determining these valuable features are contained in a highly-repetitive and complex genome almost twice the size of that in humans.

With the advent of next-generation sequencing (NGS), the barley community envisioned the sequencing of a complete barley genome (3). Chromosome sorting by flow cytometry reduced the genome complexity allowing the application of NGS to barley chromosome arms (4), which was applied to assemble the detected genes in a synteny-based virtual linear order (5). In 2012, The International Barley Sequencing Consortium (IBSC) released an extensive genome sequence resource that integrated annotated whole-genome shotgun (WGS) sequences within a physical and genetic framework (6). Bacterial artificial chromosome (BAC) and BAC-end sequences assisted the incorporation of WGS sequence data into the physical map, and the integration of the physical and genetic maps. Most of the 6,278 sequenced BACs that were included in that work were gene-bearing BACs identified from the Yu et al. (2000) library of cv. Morex (6). Subsequently, additional anchoring of WGS contigs by POPSEQ [POPulation SEQuencing (7)] led to an improved coupling of the barley physical map (8) to the genetic map. However, resolution was still quite limited as it included only the BAC sequences that were previously published (6), providing a ‘sequence-ready’ physical map (8).

Sequencing the entire barley genome has been a challenge due to difficulties resolving the abundant and complex repetitive regions during assembly (9). However, the makeup of the barley genome presents some opportune portals of entry. Estimates indicate that barley gene content is similar to that of the grass model rice, even though the latter is almost 12 times smaller (6, 10). Several studies have shown that the >80% repetitive DNA is not randomly distributed across the barley genome and that there are gene-rich regions with relatively little repetitive DNA which exhibit extensive collinearity with other grasses (e.g. 11-14). A selective sequencing strategy to target gene-containing regions of the genome has provided a feasible approach to explore and characterize the genomic features of the gene-rich portion of barley, the Triticeae model genome.

In an effort that started over a decade ago, the original Yu et al. (15) BAC library constructed from cv. Morex was screened for gene-containing BACs. Here we present the development of a minimal tiling path (MTP) of 15,711 ‘gene-bearing’ BACs, summaries of annotated and map-anchored sequences of 15,622 of these clones, and facile access to this information. An earlier version of the sequence assembly for over 2,000 of these MTP BACs was released with the 2012 genome sequence resource publication (6). Here we provide much-improved sequence assemblies for those clones, along with all of the remaining MTP of gene-bearing BAC contigs. These ~1.7 Gb of gene-rich genomic sequence expand our knowledge of the characteristic features of the gene-containing regions. Furthermore, this new resource will improve the speed and precision of map-based cloning and marker development in barley and closely related species while supporting ongoing efforts in obtaining a complete reference sequence of barley.

## RESULTS AND DISCUSSION

### A physical map of the gene-containing portion of the barley genome

The barley (cv. Morex) BAC library described by Yu et al. (15) has been extensively used for positional gene cloning (e.g. 16-20), comparative sequence analysis between related species (e.g. 20, 21) and physical mapping (6, 8). This library, composed of 313,344 BAC clones representing 6.3x haploid genome coverage, was screened with genic probes to identify a subset of 83,831 ‘gene-bearing’ BACs (hereafter referred to simply as gene-bearing BACs or GB-BACs). Rearrays encompassing these 83,831 BACs were fingerprinted using the high information contig fingerprinting (HICF) method of Luo et al. (22). Among the 72,052 clones that were effectively fingerprinted, 61,454 were assembled into 10,794 contigs (Dataset S1) using a compartmentalized assembly method (23). This assembly thus had an average of 5.7 BACs per fingerprinted contig (FPC) along with 10,598 singletons. The assembly is available at http://phymap.ucdavis.edu/barley/ in the database ‘Barley Compartmentalized PhyMap V14’.

MTP clones were chosen using the FMTP method (24) to reduce redundancy prior to sequencing. The set of MTP clones included a total of 15,711 unique BACs. Dataset S1 provides a summary of the origination of the full list of GB-BACs and these MTP clones.

### Gene-bearing MTP BAC-clone sequencing and assembly

To sequence the 15,711 unique BAC clones comprising the gene-bearing MTP we followed essentially the pooled-clone combinatorial sequencing method described by Lonardi et al. (25). In brief, eight sets of BAC clones (sets HV3 to HV10) were sequenced using Illumina HiSeq2000. High-quality reads were assigned (or *deconvoluted*) to individual BAC clones and then assembled BAC-by-BAC using Velvet v. 1.2.09 (26). Only *nodes* (i.e., Velvet contigs) with a size of at least 200 bp were used for further analysis in the present work. Assembly statistics for nodes of different minimum sizes are reported in Table S2.

A total of 15,622 BAC assemblies were obtained, which represents 99.4% of all MTP BACs attempted to be sequenced. These BAC assemblies had an average N50 of 23.9 kb and an average L50 of 2.8 nodes (Table 1). Altogether, the assembly generated 1.7 Gb of gene-rich genomic sequence, which amounts to ~33.3% of the barley genome [ca 5.1 Gb (6)] (Table 1).

**Table 1.** Statistics of the gene-bearing BAC sequence assembly for nodes ≥ 200 bp in size.

| **Chr.**<br>**arm** | **# BACs** | **Avg. # Nodes**<br>**per BAC** | **Length of assembled**<br>**reads (bp)** | **Avg. BAC**<br>**length (bp)** | **Avg.**<br>**N50** | **Avg.**<br>**L50** | **# Unique HC**<br>**gene models \*** | **# Unique LC**<br>**gene models \*** |
| --- | --- | --- | --- | --- | --- | --- | --- | --- |
| 1H | 1,959 | 19.5 | 209,800,854 | 107,096 | 23,697 | 2.8 | 2,866 | 3,502 |
| 2HS | 1,048 | 18.5 | 111,465,824 | 106,361 | 24,703 | 2.7 | 1,486 | 1,935 |
| 2HL | 1,391 | 19.1 | 149,179,441 | 107,246 | 22,978 | 2.7 | 2,241 | 2,529 |
| 3HS | 862 | 18.2 | 92,368,717 | 107,156 | 26,301 | 2.6 | 1,242 | 1,704 |
| 3HL | 1,389 | 18.8 | 148,465,120 | 106,886 | 24,223 | 2.7 | 2,132 | 2,576 |
| 4HS | 862 | 17.9 | 94,522,065 | 109,654 | 26,577 | 2.6 | 1,048 | 1,442 |
| 4HC | 60 | 15.8 | 5,866,127 | 97,769 | 28,999 | 2.2 | 20 | 40 |
| 4HL | 1,100 | 18.7 | 120,519,518 | 109,563 | 25,756 | 2.7 | 1,536 | 1,740 |
| 5HS | 640 | 19.1 | 69,074,495 | 107,929 | 24,696 | 2.8 | 812 | 1,207 |
| 5HL | 1,623 | 20.1 | 173,282,123 | 106,767 | 22,285 | 2.9 | 2,777 | 3,288 |
| 6HS | 823 | 19.7 | 87,025,477 | 105,742 | 22,248 | 2.8 | 1,070 | 1,624 |
| 6HL | 1,113 | 19.3 | 120,662,552 | 108,412 | 23,922 | 2.8 | 1,610 | 1,942 |
| 7HS | 1,196 | 19 | 129,484,046 | 108,264 | 24,580 | 2.7 | 1,770 | 2,496 |
| 7HL | 1,150 | 20.1 | 122,082,348 | 106,159 | 23,221 | 2.8 | 1,734 | 2,182 |
| NA | 406 | 41.6 | 63,956,654 | 157,529 | 16,942 | 5.2 | 994 | 1,287 |
| **All** | **15,622** | **19.7** | **1,697,755,361** | **108,677** | **23,906** | **2.8** | **15,707** | **19,330** |

\* Gene models hitting 10 or more BACs are not included in the count.

BACs were assigned to barley chromosome arms using a tool called CLARK [CLAssifier based on Reduced K-mers (27)]. Using flow-sorted materials (28) that were shotgun sequenced and assembled (see Materials and Methods), this approach allocated 15,216 BAC assemblies (97.4% of those sequenced) with high confidence to chromosome 1H or arms of chromosomes 2H-7H (Table 1; Dataset S1). The number of BACs per chromosome ranged from 1,936 for 6H to 2,439 for 2H (Table 1). We observed a linear relationship between the number of sequenced BACs per arm and the molecular size of the corresponding barley chromosome arm reported by Suchankova et al. (28) (*r*=0.953; Fig. S1). This outcome indicates that chromosomal gene content in barley is proportional to size. It should be noted that 60 BACs are located in a region that is overlapped by 4HS and 4HL cytogenetic stocks, which we defined as ‘4HC’. Some of them could be centromere clones (see below). Only 406 BACs could not be assigned to 1H or an arm of any other chromosome. Physical chimerism or cross-contamination of cultures or DNA samples during handling seem to be the most likely explanations of assignment failure for these clones, which as a group have anomalous metrics (see Table 1).

Previously predicted barley genes (6) classified into high-confidence (HC) and low-confidence (LC) were used to annotate BAC assemblies. A total of 17,386 HC and 21,175 LC gene models were found in our BAC sequences. We noticed that some gene models generated BLAST alignments to a large number of BACs, so for several calculations we excluded any gene model hitting ten or more BACs (1,679 HC and 1,845 LC genes; Dataset S2) since their inclusion would obscure more meaningful information. These gene models most often seem to contain transposable element (TE)-related sequences. Over 88% of the MTP BACs (13,809 BACs) contained at least one gene model, which indicates that the number of false positives occurring during the identification of gene-bearing BACs was low.

Our sequencing and assembly methods were validated by comparing our assemblies to those from 1,012 of the same BACs that had been previously sequenced by other institutions using 454/Roche technology (6). Very similar gene densities and gene contents were found in assemblies produced independently by these two approaches (Table 2).

**Table 2.** Comparison between gene models found in IBSC* and UCR sequence assemblies of 1,012 BACs.

| Sequencing institution | Avg. HC gene models / BAC | Total unique HC gene models | Avg. LC gene models / BAC | Total unique LC gene models |
| --- | --- | --- | --- | --- |
| IBSC* | 2.94 | 2,620 | 3.26 | 2,979 |
| UCR | 2.88 | 2,599 | 3.18 | 2,934 |
| Intersection | 2.80 | 2,502 | 3.05 | 2,794 |

High-confidence (HC) and low-confidence (LC) gene models predicted by IBSC (6) were considered. A minimum sequence length of 200 bp and an e-value of $1e^{-20}$ were the cutoffs used for the BLAST alignments. Numbers do not include gene models hitting $\geq 10$ BACs. (\*) Sequencing institutions for 454 sequencing included IPK Gatersleben, Fritz-Lipman Institute in Jena and Eurofins Scientific, and were published in IBSC (6).

### Distribution of genes in the barley genome and its correspondence with rice syntenic regions

Most of the sequenced BACs (78%) were plotted across the barley genome based on the physical coordinates provided by IBSC (6). The expected enrichment of clones towards the ends of the chromosomes was observed (Fig. 1; Fig. S2); it has been previously reported that distal regions of barley chromosomes have higher gene density than regions nearer to centromeres (6). The sequences from these BACs allowed us to explore their gene content. On average we found 2.38 HC gene models per BAC, but some BACs did not contain any previously identified gene, while others contained over 10 HC genes. We further explored the distribution of BACs containing different numbers of genes along each chromosome. As shown in Fig. 1 and Fig. S2, the location of BACs highly enriched with genes is clearly biased towards the distal ends of the chromosomes, but additional hotspots also exist (e.g. regions indicated by arrows in Fig 1). Conversely, the frequency of BACs containing zero or only one HC gene is lower toward the telomeric ends (Fig. 1, Fig S2). Peaks of BACs containing zero genes are usually located in more central positions of the chromosomes. We note that BACs in the ‘gene-bearing’ BAC list that have zero genes may have been false positives in the subjective scoring of library filter hybridizations. The uneven distribution of BACs containing at least three genes supports the idea of gene clustering that has been previously suggested for barley and other grass genomes (29-32). In contrast with the variable BAC gene-content, a uniform GC content was detected along all barley chromosomes, with chromosome averages ranging from 44.4% (3H and 7H) to 44.6% (4H) and an average GC content for all BACs of 44.5% (SD = 1.3%). This constant GC content was also found in wheat BACs located in different regions of 3B (30) and in 1BS BAC-end sequences (32). The BAC GC-content percentages that we observe are similar to the GC composition of previously studied barley gene-bearing BACs (17, 20) and comparable to those of rye (33) and wheat genomes (30, 32). Although our dataset is biased toward gene-containing BACs, we did not find any significant difference in GC content between gene-rich and gene-poor BACs.

**Fig. 1.**
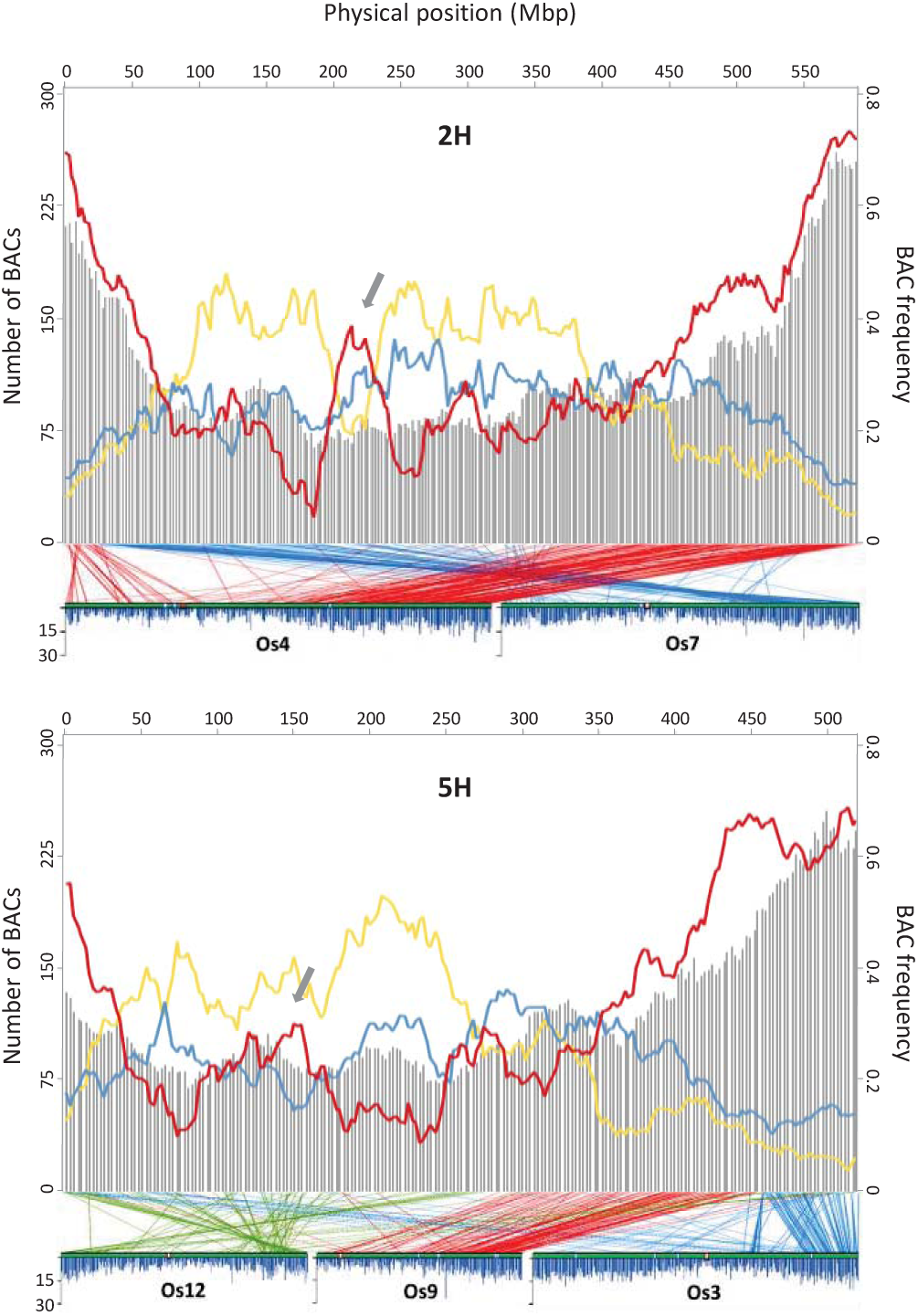
BAC distribution along barley chromosomes 2H and 5H and syntenic relationships with rice chromosomes. Grey bars represent the number of sequenced barley BACs and their units are shown on the left Y-axis. Colored lines represent the proportion of BACs containing only 1 HC gene model (blue), 3 or more HC genes (red) or 0 HC gene models (yellow), and the scale is shown on the right Y-axis. BAC densities are calculated for a sliding window of 40 Mb at 2.5 Mb intervals based on the physical coordinates (archived golden path) provided by IBSC (6). Gray arrows indicate gene-dense regions different from distal ends. Barley-rice synteny is represented by lines connecting each mapped BAC to the position on the rice genome determined by BLASTX (see Materials and Methods). Densities of expressed rice genes across chromosomes are also displayed [adapted from Supplementary Figure 2 in (10)], where blue bars indicate the frequency of gene models in 100 kb windows, red boxes indicate centromeres and white boxes represent physical gaps.

To explore barley-rice synteny, each barley BAC DNA sequence was compared to rice translated gene models available at the Rice Genome Annotation Project database (http://rice.plantbiology.msu.edu/) using BLASTX (see Materials and Methods). The syntenic relationships between rice and barley that were revealed (Fig. 1 and Fig. S2, lower plots) are consistent with previously reported observations (5, 34, 35). Single rice chromosomes (Os1 and Os2) are largely syntenic with barley 3H and 6H, respectively. Barley 5H has a more complex synteny with rice, sharing major syntenic regions with at least three rice chromosomes (Os3, Os9 and Os12). Each of the remaining barley chromosomes shares major blocks of conserved synteny with two rice chromosomes. In terms of gene content, we observed that the barley gene-enriched chromosome regions tend to correspond to rice regions of high gene density [obtained from (10)]. A clear example is barley chromosome 5H, where the gene-dense portion extends further from the long-arm distal end than in any other chromosome. This is the only distal region sharing synteny with gene-enriched distal portions of two rice chromosomes. Indeed, the 5HL distal region shows two peaks of gene enrichment, which coincide with each of the two ancestors of rice chromosomes (Fig. 1). In sharp contrast, the prominent peak that is visible in the central region of 2H does not have clear synteny to a gene-rich region of the rice genome (Fig. 1).

### Centromeric region of 4H

As mentioned above, 60 BACs were assigned to the region of overlap between the long and short flow-sorted arms of chromosome 4H (4HC), some of which may be centromere clones. Only two of those BACs (0143O21 and 0474D04) contained a mapped OPA SNP [1_0424, (35)], which was previously confirmed in the pericentromeric region of 4H (36). We also explored the gene content of these BACs. Only 14 BACs contain HC gene models when excluding highly frequent gene models (mostly TE-related) (Table S2). This is an expected finding since centromeric regions are known to have low gene density. Although we did not find any annotated gene encoding the highly conserved centromere-specific histone cenH3 (37), we found genes previously identified in the centromere of rice chromosome 3 [ribosomal protein S5 and magnesium chelatase (38)] and a gene for retinoblastoma-related protein, which has been recently shown to play an essential regulatory role in the assembly of cenH3 at Arabidopsis centromeres (39). Satellite sequences and centromere-specific retrotransposons are the major constituents of centromeres in plants. While satellite repeats are problematic due to difficulties in the assembly of these tandem repeats, several retrotransposons including the conserved Ty3/gypsy type (40, 41) were found when inspecting highly frequent genes in these BACs (Table S2). These observations may provide useful leads for further studies of the centromere of barley chromosome 4H.

Although specific *k*-mers were identified for centromeric regions of overlap of all other barley chromosomes except 1H (27), none of our sequenced BACs were assigned to those regions. This is probably because, in 4H, the region shared between flow-sorted short and long arms is larger than that in any other chromosome. However, when additional barley BACs are sequenced, it may be possible to identify centromeric BACs for other barley chromosomes. The use of analysis tools based on discriminative *k*-mer (such as CLARK) in combination with chromosome arm sequence data could be used as an approach to define centromeric regions in other species where flow-sorted arms exist (i.e. bread wheat).

### Relationship between gene density and recombination rate

Triticeae chromosomes exhibit an increase in gene density and recombination rate along the centromere-telomere axis (42). This general trend has been observed in sequence data from barley (6, 43), wheat (32, 44), and *Ae. tauschii* (45), and it is common to other grasses (i.e. Brachypodium, rice, and maize). The possibility of examining larger pieces of genome sequence (i.e., BACs) accounting for one third of the barley genome allowed us to take a closer look at how recombination rate distribution relates to gene distribution. As shown in Fig. 2, recombination frequency generally increases along the centromere-telomere axis. Similarly, gene density distribution is not uniform along the chromosomes and it is usually correlated with recombination frequency (i.e., higher in distal ends). However, we also observe regions that clearly deviate from these general characteristics; there are regions with relatively high gene density embedded within areas with suppressed recombination. The two clearest examples are in chromosomes 2H and 5H (Fig. 2, grey arrows). The 2H region coincides with the prominent peak of gene-rich BACs shown in Fig. 1 and comprises ~18 Mb. A total of 50 BACs containing 84 HC unique gene models (Dataset S3) are located in this region of 2H. GO-term enrichment analyses performed in agriGO (46) using GO terms available from MIPS [http://pgsb.helmholtz-muenchen.de/plant/barley/, (47)] revealed a slight enrichment for genes belonging to category ‘cellular component organization’, including alpha-tubulins, peptide chain release factors, and a thylakoid-formation protein (thf1) (Dataset S3), which perform essential cellular functions. This barley region shares conserved synteny with the *Ae. tauschii* 2D region extending from 168 to 182 Mb in the Luo et al. (45) physical map (ctg3995, ctg157 and ctg3482).

**Fig. 2.**
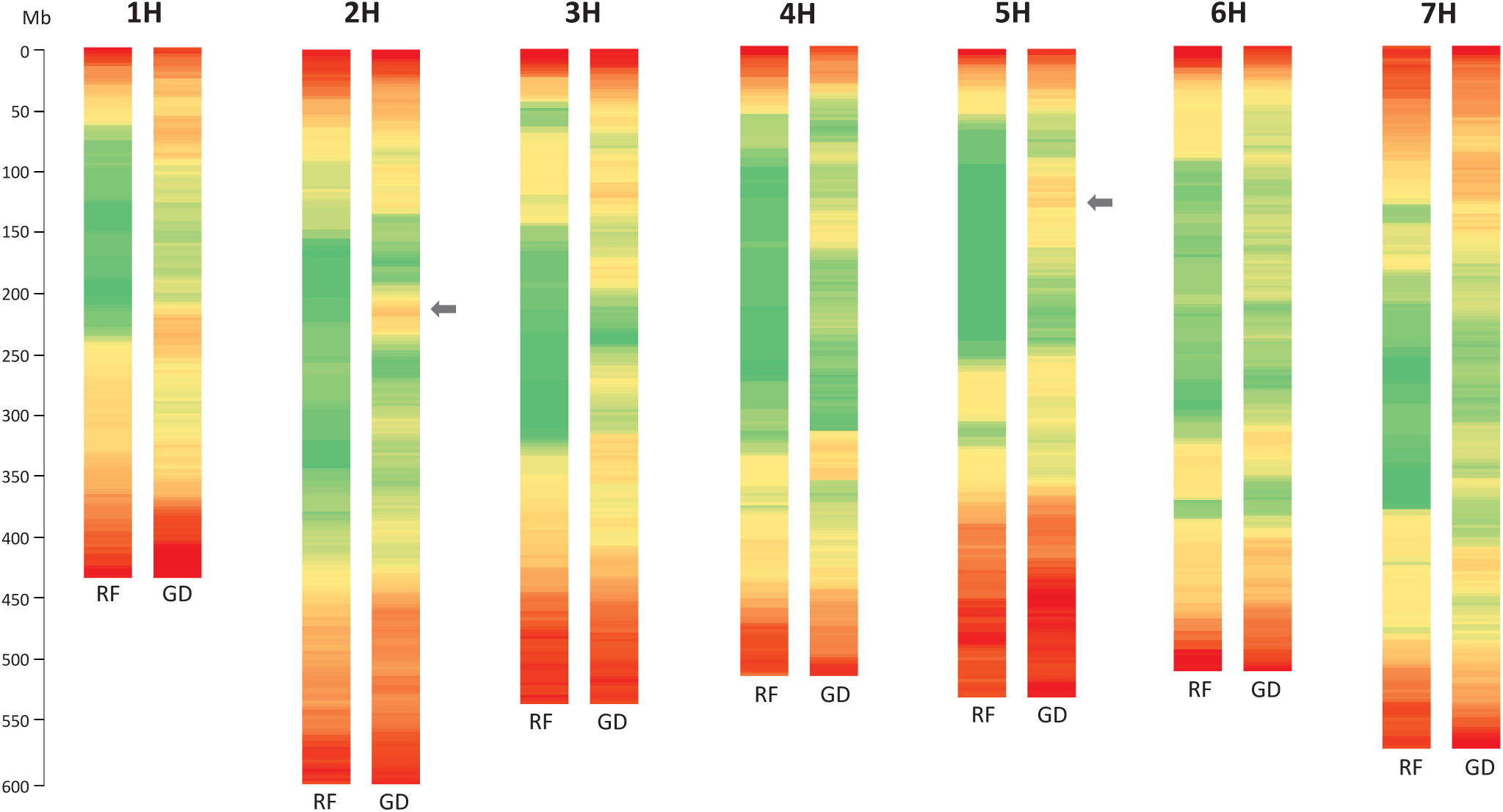
Relationship between recombination frequency (RF) and gene density (GD) along the seven barley chromosomes. Recombination rates are calculated from the cM/Mb ratios in sliding windows of 40 Mb with 2.5 Mb increments, and are represented by a color gradient from green (RF=0) to red (RF=1.14). Gene densities are estimated based on the total number of unique HC genes per window with respect to the total number of sequenced BACs assigned to that window, and are represented by the same color gradient from green (GD=0.67) to red (GD=3.16). Grey arrows indicate most evident genomic regions of relatively high gene density and very low recombination.

The 5H region is much larger, extending for approximately 60 Mb, including 136 BACs and 174 HC gene models. Interestingly, the GO category ‘response to biotic stimulus’ was a highly overrepresented. A set of seven genes belonging to the ‘Bet v I type allergen’ family of proteins is responsible for this enrichment. Different pollen allergens and pathogenesis-related proteins (i.e. STH-21; Dataset S3) are included in this family. Due to their homology with pathogenesis-related proteins, Bet v I pollen allergens are considered to be involved in pathogen resistance of pollen (48). This recombination cold-spot did not contain any of the rapidly evolving nucleotide binding, leucine-rich-repeat (NLR) encoding R genes, which tend to cluster in high-recombination distal regions of the barley chromosomes (6). The other enriched GO category in the highlighted 5H region was ‘DNA replication’, involving genes with more conserved functions. Some barley BACs in this region are syntenic to *Ae. tauschii* sequences at 94 Mb on 5D [ctg582, (45)].

A list of HC genes located in the aforementioned regions is provided in Dataset S3. To facilitate the exploration of these and other regions of the barley genome, Dataset S4 contains the recombination frequency and gene density data corresponding to Fig. 2 with detailed information of barley physical and genetic positions. As recombination determines the extent of linkage disequilibrium (LD), it impacts both genome-wide association studies (GWAS) and genomic selection (GS). Regions of high LD result in spurious marker-trait associations in GWAS. Modification of GS models to emphasize recombinants in low-recombination regions can be particularly important in regions with higher gene content where genetic load may also be higher (49). Similarly, marker-assisted trait introgression and map-based cloning are affected by recombination, since large populations may be required to break linkage drag or to find markers within a short physical distance of the target gene. Thus, knowledge of recombination rates and gene densities provided in this work is of relevance for the effective use of genetic variation in barley and related species.

### **HarvEST:Barley allows easy access to GB-BAC sequence assemblies and synteny with *Ae. tauschii***

To successfully deploy genome sequence information for crop improvement, it is critical to make it easily accessible. We have developed an online interface to provide access to the latest sequence assemblies (v.4.1) from 15,622 MTP gene-bearing BACs (http://harvest-web.org/hweb/utilmenu.wc). BAC sequences and their annotations can be retrieved by ‘BAC address’, which can be obtained by BLAST via http://www.harvest-blast.org/ using the user’s input sequence. Since high-throughput SNP genotyping (35, 50) is routinely used in modern barley breeding, genome data can also be downloaded using a ‘SNP name’ as a query. Alternatively, HarvEST:Barley (http://harvest.ucr.edu/) may be installed on a personal computer to export BAC sequences, gene annotations and both physical and genetic map coordinates in a similar manner, with additional options of exporting BACs by chromosome arm or genetic map interval. HarvEST:Barley is available for Microsoft Windows and does not require an internet connection to operate. These two interfaces also provide the option of accessing information from only the specified BACs or all BACs in the same contig.

Along with the 15,622 BAC sequences described in the present work, the databases noted above contain sequences from additional BACs from the Yu et al. (15) library. This includes: 3,153 BAC assemblies generated using 454 sequencing (6); 50 BACs fully sequenced at the Joint Genome Institute by Sanger sequencing; and 21 BAC sequences available from the National Center for Bioinformatics (NCBI) database. A list of these BACs sequenced by other methods, some of which coincide with the BACs that we sequenced, can be found in Dataset S5.

The latest version of HarvEST:Barley has implemented a barley-*Aegilops tauschii* synteny viewer to assist comparative genomic studies among Triticeae species. This is in addition to the previously available barley-rice and barley-*Brachypodium distachyon* synteny displays, all of which can be retrieved by selecting either of the two latest barley genetic consensus maps (1, 36).

## MATERIALS AND METHODS

### Identification of gene-containing BACs

A 6.3x haploid-genome-equivalent barley BAC library [(15), http://www.genome.clemson.edu] was obtained as library filter sets from Clemson University Genomics Institute, as were rearrayed cultures of BAC clones used for fingerprinting. This library was constructed using *Hind*III partially digested cultivar Morex DNA ligated to pBeloBAC11 vector and arrayed on 17 filters, each with 18,432 clones. Compilation of 81,831 gene-bearing BACs was accomplished in two ways. A total of 21,689 BACs were identified from information provided by several researchers based on their prior work (Dataset S1). In addition, oligonucleotide probes (‘overgos’) designed mainly from transcript sequences (EST unigenes) were used in hybridizations to identify 72,141 putative gene-bearing BACs. The union of these two sets was 83,831 unique GB-BACs. Information regarding the pools of probes used for BAC identification and details of the hybridization process can be found in Supporting Materials and Methods.

As the hybridizations using pools of overgos progressed, we observed gradually fewer newly identified BACs per hybridization. The frequency of new BACs identified for each of the large pools of probes generally diminished from a starting range of 60-85% to a range of 27- 57%. To provide an estimate of the percentage of all possible gene-bearing BACs identified in this work, the hybridization data were randomly shuffled and sampled 10,000 times to plot the number of unique BACs identified as a function of the number of probe pools applied. Extrapolation provided an estimate of the number of pools that would be necessary to approach the asymptotic limit of the number of gene-bearing BACs. From this treatment of the data, we estimated 107,882 gene-bearing BACs in the 6.3X Morex library (Fig. S3), which is roughly 1/3 of all BACs genome-wide. However, the nature of our BAC detection method (genic probes) made it more likely that we would find a BAC containing multiple genes than a BAC carrying only one gene, and as a consequence the asymptotic limit could be higher than indicated by this extrapolation.

### BAC-clone fingerprinting and assembly

BAC DNA was isolated and fingerprinted according to Luo et al. (22). BAC clones maintained in a 384-well plate were inoculated into four 96-deep well plates containing 1.2 mL 2X YT medium with 12.5 μg/mL chloramphenicol and grown at 37° C for 24 h. BAC DNA was isolated with the Qiagen R.E.A.L 96-Prep kit either manually or by Qiagen robot. 0.5-1.2 μg of DNA was then digested with 5 units each of *BamH*I, *EcoR*I, *Xba*I, *Xho*I and *Hae*III enzymes and transferred to SNaPshot multiplex labeling solution. Capillary electrophoresis was performed on the DNA using ABI internal size standard LIZ-500 (35-500 bp) in an ABI 3730 DNA sequencer (Applied Biosystems, USA). GenoProfiler software (51) was used for automated editing of sized fingerprinting profiles generated by the ABI Genetic Analyzers (Applied Biosystems, USA). The batch-processing module extracts sized fragment information either directly from the ABI raw trace files or from data files exported from GeneMapper (Applied Biosystems, USA) or other size calling software, removes background noise and undesired fragments, and generates fragment size files. High-throughput fingerprinting was attempted for 83,831 clones. 2,967 clones (5 %) were deleted during fingerprint editing due to lack of insert, failed fingerprinting, or being ignored by the GenoProfiler software. Clones containing four or fewer fragments in the range of 50 to 500 bp) provided insufficient information to be included in contig assembly. The clones from this library had an average of 92 fragments per clone in the sized range.

BAC clones were assembled into contigs using a compartmentalized approach (23). Each group of BACs identified by each probe pool was first assembled independently using Fingerprint Contigs (FPC) software v8.1 (52, 53). Contig merging and redundancy removal steps were then applied (23) to finalize the assembly. Key elements of this compartmentalized process can be found in Supporting Materials and Methods.

### Construction of minimal tiling paths

Two sets of MTP clones were computed. The first one contained 2,638 BACs from data that were available relatively early in this work, including the majority of GB-BACs provided from prior work and GB-BACs containing abiotic-stress regulated genes and others targeted by a SNP genotyping assay referred to as POPA1 (35). These BACs are referred to later as MTP subsets 1-3. Subsequently, after the full list of 83,831 GB-BACs had been compiled, the second set of MTP clones, 13,182 BACs, was developed. BACs from this second MTP set, which generally avoided duplication with the first set of MTP clones, are referred to later as MTP subsets 4-9.

### MTP sequencing and assembly

MTP BACs were paired-end sequenced (2 x 100 bases) using Illumina HiSeq2000 (Illumina, Inc, San Diego, CA, USA). Sequencing was done in eight sets of BAC clones (HV3-10) applying a combinatorial pooling design (25). In brief, this approach takes advantage of the current high-throughput sequencing instruments to *de novo* sequence thousands of BAC clones in pools that are designed to identify each BAC within the pooling pattern, hence avoiding exhaustive DNA barcoding. Reads in each pool (after quality trimming) were ‘sliced’ into smaller samples of optimal size as explained in detail by Lonardi et al. (54), deconvoluted, and then assembled BAC-by-BAC using Velvet v.1.2.09 (26). Assembly of the BACs with SPAdes or IDBA (56) did not provide a clear objective improvement in assembly quality. Slicing the data into subsets was a key step to improving the quality of the assemblies. A threshold cutoff of 35,000 high-quality reads was applied to consider a BAC as ‘sequenced’. Statistics for assembly were collected using the Assemblathon script (K. Bradnam, Genome Center, UC Davis). The BAC sequence reads are available in NCBI under accession numbers SRX143098, SRX138629, SRX104477, SRX119494, SRX142443, SRX142540, SRX142565, SRX142566 and SRX268475.

High-confidence (HC) and low-confidence (LC) gene models predicted by IBSC (6) were used to annotate the BAC assemblies, using a minimum sequence length of 200 bp and an e-value of 1e^-20^as the cutoffs for the BLAST alignments. We ignored any gene model hitting at least 10 BACs for most analyses.

### Chromosome-arm assignment of BACs

We used CLARK, a supervised classification method, to assign BACs to chromosome arms (27). Briefly, CLARK can accurately classify ‘objects’ (e.g., BACs) to ‘targets’ (e.g., chromosome arms) by reducing the problem to a *k*-mer comparison of the corresponding sequences. CLARK differs from other *k*-mer based methods because it considers only *k*-mers that are specific (or discriminative) to each target. It does so in the preprocessing phase by discarding any *k*-mer that appears in two or more targets, except in the case of *k*-mers shared by only both arms of the same chromosome, which are used to define ‘centromeric’ regions of overlap. Additionally, CLARK disregards very rare *k*-mers, which tend to be spurious from sequencing errors. Using *k*=19 and by discarding 19-mers that appeared only once (27), we have accepted only assignments with confidence scores > 0.75. ‘Targets’ were reads generated by Illumina whole-genome shotgun sequencing of barley flow-sorted chromosome 1H and arms of chromosomes 2H-7H that were assembled using SOAPdenovo (57). The chromosomes were purified by flow cytometric sorting as described by Suchánková et al. (28) and their DNA amplified following the procedure of Šimková et al. (58). Flow-sorted chromosome arm sequences can be found at NCBI under accession no. SRX143974.

### Validation of the sequence assembly

A total of 1,037 gene-bearing BACs from the Yu et al. library (15) were previously sequenced (454 Life Sciences technology) and assembled by other institutions, and included in the barley genome sequence resource (6). Our sequence assemblies for 1,037 BACs of the same address BACs were compared using MUMmer 3.0 (59). We removed 25 BACs which had <33% alignment with each other, attributing this level of disagreement to rearray errors, extensive cross-contamination or extreme instability of a BAC in the *Escherichia coli* host. The remaining 1,012 BACs that were sequenced independently in the present work and previously were blasted against HC and LC gene models predicted by IBSC (6), using a minimum sequence length of 200 bp and an e-value of 1e^-20^. Gene models found in our sequence assemblies and in IBSC were compared, after excluding gene models hitting at least 10 BACs (Table 2).

### Synteny analysis

For barley-rice synteny, each barley BAC DNA sequence was compared to rice translated gene models available at the Rice Genome Annotation Project database (http://rice.plantbiology.msu.edu/). All BLASTX hits with an e-value of −20 or better were tallied for each BAC. The rice chromosome with the plurality of matches was then taken as the correct rice chromosome. A mean value of chromosome coordinates was then calculated for the matched rice gene models on this chromosome to assign a rice chromosome position to each BAC. A rice genome position was then assigned to each entire BAC contig by a similar voting method, accepting the plurality of rice chromosomes for the contig and the mean value within the matching rice chromosome as the position of the BACs in this contig. These rice genome coordinates were then used to align the seven barley chromosomes with the twelve rice chromosomes (Fig. 1 and Fig S2).

For barley-*Aegilops tauschii* synteny, each SNP design sequence for the iSelect genotyping assay (45), downloaded from http://probes.pw.usda.gov/WheatDMarker/downloads/, was matched by BLAST to the extended genome sequences that were also available from http://probes.pw.usda.gov/WheatDMarker/downloads/. The linkage group and cM coordinates for each SNP marker published in Luo et al. (45) were then associated with each wheat D genome iSelect SNP assay design sequence. These wheat SNP design sequences were matched by BLAST to the sequences of barley BACs described in the present work, many of which contained sequences matching barley SNP assay design sequences (35, 50). The arm assignment for each barley BAC was taken from Dataset S1, to limit the BACs to those where the barley and wheat SNPs mapped to orthologous chromosomes. The barley-wheat D synteny viewer in HarvEST:Barley (for Windows) is based on these relationships.

## ACKNOWLEDGEMENTS

This work was supported by the USDA Initiative for Future Agriculture and Food Systems 01- 52100-11346, North American Barley Genome Project (USDA-CSREES 2001-34213-10511), USDA-CSREES National Research Initiative (NRI) 2002-35300-12548, NSF Plant Genome Research Program DBI-0321756, BarleyCAP (USDA-CSREES-NRI 2006-55606-16722 and USDA-AFRI-NIFA 2009-85606-05701), USDA-AFRI-NIFA 2009-65300-05645, TriticeaeCAP (USDA-NIFA 2010-15718-10), NSF-ABI DBI-1062301, and UC Riverside Agricultural Experiment Station Hatch Project CA-R-BPS-5306-H. The work conducted by the U.S. Department of Energy Joint Genome Institute was supported by the Office of Science of the U.S. Department of Energy under Contract No. DE-AC02-05CH11231. H.Š and J.D. have been supported by grant award LO1204 from the National Program of Sustainability I. The authors also thank the following individuals: for communications regarding gene-bearing BAC IDs (Nick Collins, Jorge Dubcovsky, Katherine Feuillet, Kulvinder Gill, Yong Gu, David Laurie, Saghai Maroof, Tim Sutton, Pingsha Hu); for BAC clone identification (Faith Lin, Ginger Mok, Hung Le); for provision of fee-for-service Morex barley research materials (Michael Atkins, Clemson University Genomics Institute), for provision of fee-for-service DNA sequencing (John Weger, UC Riverside Genomics Core Facility), and for other technical assistance (Gianfranco Ciardo, Raymond Fenton, Hung Le, Harkamal Walia).

